# Synaptosome microRNAs regulate synapse functions in Alzheimer’s disease

**DOI:** 10.1101/2021.12.15.472852

**Authors:** Subodh Kumar, Erika Orlov, Prashanth Gowda, Chhanda Bose, Russell H. Swerdlow, Debomoy K. Lahiri, P. Hemachandra Reddy

**Author notes:** **Address for correspondence** Subodh Kumar, Ph.D., Internal Medicine Department, Texas Tech University Health Sciences Center, 3601 4th Street, Lubbock TX 79430, United States. **Co-correspondence** P. Hemachandra Reddy, Ph.D., Professor of Internal Medicine, Cell Biology & Biochemistry, Neuroscience & Pharmacology, Neurology, Public Health and School of Health Professions, Texas Tech University Health Sciences Center, 3601 4th Street, Lubbock TX 79430, United States.

## Abstract

MicroRNAs (miRNAs) are found in nerve terminals, synaptic vesicles, and synaptosomes, but it is unclear whether synaptic and cytosolic miRNA populations differ in Alzheimer’s disease (AD) or if synaptosomal miRNAs affect AD synapse activity. To address these questions, we generated synaptosomes and cytosolic fractions from postmortem brains of AD and unaffected control (UC) samples and analyzed them using a global Affymetrix miRNAs microarray platform. A group of miRNAs significantly differed (p<0.0001) with high fold changes variance (+/- >200-fold) in their expressions in different comparisons- 1) UC synaptosome vs UC cytosol, 2) AD synaptosomes vs AD cytosol, 3) AD cytosol vs UC cytosol, and 4) AD synaptosomes vs UC synaptosomes. MiRNAs data analysis revealed that some potential miRNAs were consistently different across sample groups. These differentially expressed miRNAs were further validated using AD postmortem brains, brains of APP transgenic (Tg2576), Tau transgenic (P301L), and wild type mice. The miR-501-3p, miR-502-3p and miR-877-5p were identified as potential synaptosomal miRNAs upregulated with disease progression based on AD Braak stages. Gene Ontology Enrichment and Ingenuity Pathway Analysis of synaptosomal miRNAs showed the involvement of miRNAs in nervous system development, cell junction organization, synapse assembly formation, and function of GABAergic synapse. This is the first description of synaptic versus cytosolic miRNAs in AD and their significance in synapse function.

## Introduction

Alzheimer’s disease (AD) progresses with synaptic failure caused by amyloid beta (Aβ) and phosphorylated tau (p-tau) toxicities at synapses. In aged individuals, the numbers of AD cases are increasing gradually, and by mid-century, the number of Americans age, 65 and older with Alzheimer’s dementia may grow to 13.8 million (Alzheimer’s disease facts and figures 2021). This represents a steep increase from the estimated 5.8 million Americans age, 65 and older who have Alzheimer’s dementia today.

Synaptic dysfunction or poor pre-synaptic and post-synaptic activities leads to the synaptic degeneration and neuron death in AD (Forner et al. 2017; Marsh and Alifragis 2018; Kashyap et al. 2019). It is well known that synapse loss and dysfunction are the main physiological and pathological hallmarks of AD (Selkoe 2002; Chen et al. 2019; Ahmad and Liu 2020; Colom-Cadena et al. 2020).

Synapses are the key components for healthy brain functioning. Synapse integrity (number, structure and functions) are crucial for a balanced neurotransmission and to maintain healthy synaptic and cognitive functions of the brain. Synapse components can be extracted from postmortem brains in an intact form referred as ‘synaptosome or synapto-neurosomes’. Synaptosomes are the best neural cell component to study the synapse dysfunction in multiple neurodegenerative diseases, particularly in AD, where the synaptosome structure and functions are altered due to Aβ and p-tau accumulations (Kumar and Reddy 2020). During early AD progression, synapses are the first targets that are hit by Aβ and p-tau toxicities (Reddy et al. 2012; Spires-Jones and Hyman 2014; Jackson et al. 2019). Multiple synaptic events are disturbed in AD, such as axonal transport, synapse mitochondrial function, synaptic vesicle trafficking, release and cycling, alteration of Ca^++^ influx, neurotransmitter release, impaired receptors, inflammation and synaptotoxicity (Kumar and Reddy 2020; Calkins et al. 2011; Swerdlow 2020; Weidling and Swerdlow 2020; Kodavati et al. 2020; Ammal Kaidery et al. 2021; John and Reddy 2021).

MicroRNAs (miRNAs) are present throughout cells (Kumar and Reddy 2020). Some miRNAs are localized to subcellular compartments, including the rough endoplasmic reticulum, processing (P)-bodies, stress granules the trans-Golgi network, early/late endosomes, multivesicular bodies, lysosomes and mitochondria (Kumar and Reddy 2020; O’Brien et al. 2018). Several studies identified the presence of miRNAs at the synapse and in synaptosomal fractions and determined their important roles in the regulation of local protein synthesis (Lugli et al. 2008; Xu et al. 2013; Li et al. 2015; Boese et al. 2016). Even synaptic vesicles extracted from mouse central nervous system contain several small RNAs, transfer-RNAs and miRNAs (Li et al. 2015). Additionally, miRNAs were found to be abundantly expressed within synaptoneurosomes isolated from prion-infected forebrain (Boese et al. 2016).

Since the 1980s, researchers began using synaptosomes prepared from postmortem brains to study AD-associated deficits in neurotransmission, including dysfunction of excitatory synapse acetylcholine, glutamate or aspartate, and inhibitory synapse glycine or (gamma-aminobutyric acid) GABA systems (Rylett et al. 1983; Rajmohan and Reddy 2017; Lauterborn et al. 2021). A decrease in GABAergic synapse activity and inhibitory interneurons could contribute to AD progression and cognitive deficits in human and AD mouse models (Govindpani et al. 2017; Hollnagel et al. 2019; Xu et al. 2020; Jiménez-Balado and Eich 2021). Synaptic disturbances at the excitatory and inhibitory synapse in the forebrain have been found to contribute the progression of AD and dementia (Lauterborn et al. 2021). Recent synaptosomal studies have revealed decreased levels of neprilysin in AD patients (Jhou and Tai 2017). Neprilysin plays a key role in the clearance of Aβ.

Recently, it is well acknowledged that miRNAs exert widespread regulation over the translation and degradation of their target genes in nervous system (Schratt 2009; Siegel et al. 2011; Wingo et al. 2020). Increasing evidence suggests that quite a few specific miRNAs play important roles in various aspects of synaptic plasticity, including synaptic activity, synaptic development, synaptogenesis, synaptic morphology, synaptic remodeling, synaptic scaling, synaptic excitability, synaptic ATP production and synaptic integrity (Kumar and Reddy 2020; John and Reddy 2021; Reddy and Beal 2008; Smallheiser 2014; Ye et al. 2019; John et al. 2020; Gowda et al. 2021). More importantly, the miRNA-mediated regulation of synaptic plasticity is not only responsible for synapse development and function but is also involved in the pathophysiology of plasticity-related diseases including AD (John and Reddy 2021; Smallheiser 2014; Ye et al. 2019).

MiRNAs are the potential regulators of gene(s) and gene products and their therapeutic relevance have been explored in human diseases, including AD (Lahiri and Maloney 2010; Long and Lahiri 2011; Long et al. 2019; Chopra et al. 2020; Lukiw 2020; Zhao et al. 2019; Kumar and Reddy 2016). The role of miRNAs has been exposed in the regulation of synaptic activity in the case of AD (Kumar and Reddy 2020).

MiRNAs which enrich at the synapse directly regulate local protein synthesis involved in multiple synaptic functions and governing synaptic plasticity (Lugli et al. 2008; Xu et al. 2013; Li et al. 2015; Boese et al. 2016; Zolochevska and Taglialatela 2020; Yoshino et al. 2021).

However, the role of synaptosome-specific miRNAs is not determined in the progression of AD. There are no published reports about synaptosome-specific miRNAs for AD thus far.

Futhermore, it is unclear whether synaptosomal miRNAs are different from cytosolic miRNAs. Hence, this study classified synaptosomal versus cytosolic miRNAs and unfurled the possible molecular link between synaptosomal miRNAs and AD progression. Our study addressed four previously unknown important research questions- 1) Are miRNA(s) levels altered at the synaptosome in AD? 2) If so, are synapse miRNAs expressed differently in AD than in a healthy state? 3) Are synaptosomal miRNAs expressed differentially in the cytosol? and 4) What function do synaptosomal miRNAs play in synaptic activity and neurotransmission in AD? Overall, the focus of this study is to discover synaptosomal miRNAs and understand their positive and negative roles in AD progression.

## Results

### Synaptosomes preparations from postmortem brains

Increased levels of APP and Tau proteins were detected in AD cases compared to UC samples (SI Fig. 1). Next, these samples were processed for synaptosome preparation downstream applications (Fig. 1A). Fig. 1B, showed a representative immunoblot for SNAP25, synaptophysin and PSD95 and cytosolic/nuclear proteins elF1a and PCNA. Densitometry analysis showed significantly increased levels of SNAP25, synaptophysin and PSD95 in the synaptosome fraction and reduced levels in cytosolic fraction (Fig. 1C). SNAP25 and PSD95 were completely absent from the cytosolic fraction, however synaptophysin was detected in the cytosolic fraction, which was also as reported by other researchers (Scherma et al. 2020). On the other hand, elF1a and PCNA protein levels were higher in cytosol. qRT PCR analysis also showed increased expression of SNAP25, synaptophysin and PSD95 genes in the synaptosomes relative to the cytosol and reduced expressions of elF1a and PCNA in the synaptosomes fraction relative to the cytosol (Fig. 1D). These results confirm a precise separation of cytosolic and synaptosomes fractions.

**Figure 1.**
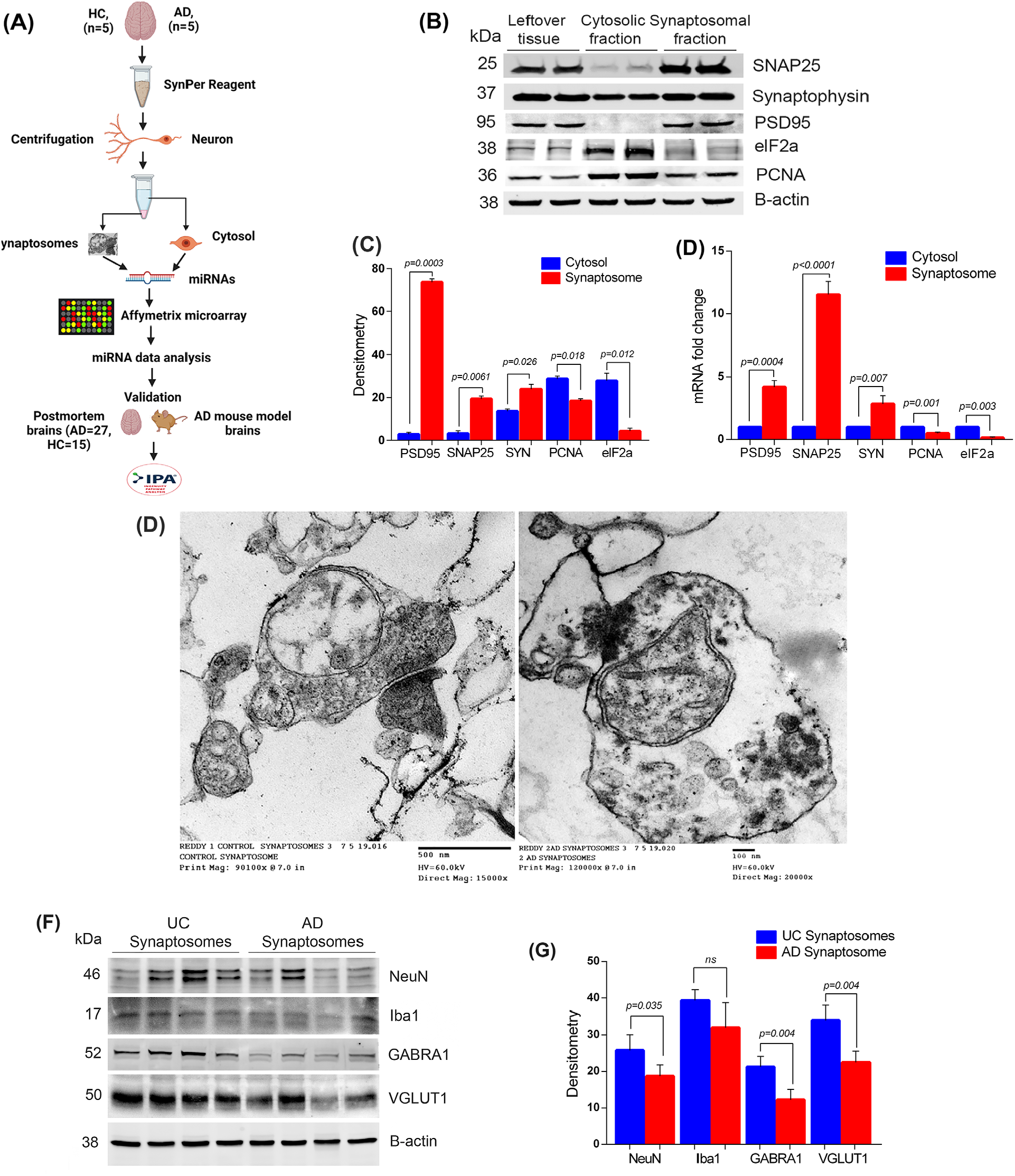
Extraction and characterization of Synaptosomes. **(A)** Brief workflow of the current study. **(B)** Immunoblotting analysis of synaptic (SNAP25, Synaptophysin and PSD95) and cytosolic (elF1a and PCNA) proteins in cytosolic fraction, synaptosomal fraction and leftover tissue debris of unaffected control postmortem brain tissues. **(C)** Densitometry analysis of synaptic and cytosolic proteins. Synaptic proteins levels (PSD95; p=0.003), (SNAP25; p=0.0061), (Synaptophysin; p=0.026) were significantly higher in synaptosomes and cytosolic proteins (elF1a; p=0.012) and (PCNA; p=0.018) levels were significantly lower in synaptosomes relative to cytosol. **(D)** qRT-PCR analysis for mRNA fold change analysis of synaptic and cytosolic genes in cytosolic and synaptosomal fractions (n=5). **(E)** TEM analysis of synapse assembly in synaptosomal fraction from unaffected control and AD patients’ postmortem brains (500 and 100 nm magnification). Electron micrograph shows synapse components mitochondria, synaptic vesicles, postsynaptic density, and synaptic cleft. **(F)** Immunoblotting analysis of brain cells markers (Neuron-NeuN; Microglia-Iba1), excitatory synapse marker (VGLUT1) and inhibitory synapse marker (GABARA1) proteins in unaffected controls (n=4) and AD (n=4) synaptosomes. **(G)** Densitometry analysis of NeuN, Iba1, VGLUT1 and GABARA1 proteins in unaffected controls and AD synaptosomes.

Next, we processed the synaptosomes fraction from AD patients and UC for TEM analysis (Fig. 1E). The electron micrograph revealed the distinct synapse assembly and intact synaptosomes with all the components-mitochondria, synaptic vesicles, endosomes, post-synaptic density protein and synaptic cleft. The mitochondrial structure and synaptic cleft were found to be distorted in AD postmortem brains, while it was intact in control samples. These results confirmed the purity and integrity of synapse and synaptosomes fraction.

Further, to confirm the brain cells specificity of synaptosomes, we checked the levels of cell type markers (NeuN- Neuron, Iba1- Microglia, GFAP- Astrocytes). We found the significantly detectable levels of NeuN and Iba1 proteins (but not GFAP) in both UC and AD synaptosomes (Fig. 1F). NeuN level was found to be significantly reduced (p=0.035) in AD synaptosmes relative to UC synaptosomes (Fig. 1G). We did not see any significant difference in Iba1 levels in AD vs UC synaptosome. These observations confirm the neuron specificity of synaptosomes.

We also characterized the synaptosomes as excitatory or inhibitory based on the levels of excitatory and inhibitory synapse markers (VGLUT1 and GABRA1). Immunoblots in Fig. 1F showed the levels of both markers in UC and AD synaptosomes. The levels of VGLUT1 (p=0.004) and GABRA1 (p=0.004) proteins were significantly reduced in AD synaptosomes relative to UC synaptosomes (Fig. 1G). These observations confirmed the presence of both types of synapses in synaptosomes fraction with their reduction in AD brains.

### MicroRNAs expression in UC synaptosomes vs UC cytosol

The miRNA microarray data of synaptosomal and cytosolic fractions were analyzed by Transcription analysis console v.4. A total of 43 mature miRNAs were found to be deregulated in UC synaptosomal fraction relative to UC cytosolic fraction (SI Table 4). As shown in SI Table 4, the 20 Homosapiens (hsa) miRNAs were highly expressed in the synaptosomes and low in the cytosol. These observations indicate that highly expressed miRNAs in synaptosomes have functional importance of synaptosomal function. The 23 hsa-miRNAs (SI Table 4) were highly expressed in the cytosol and showed reduced expression in the synaptosomes, strongly suggesting that these miRNAs have cytosolic relevance in the healthy state.

MiRNAs were characterized on several selection criteria - fold change, standard deviation, p-values, expression priority, transcript ID, chromosome location, strand specificity, start and stop codon, targeted and validated gene symbols (SI Table 4). Fig. 2A shows the hierarchical clustering and heat map of significantly deregulated miRNAs with their ID numbers. As a result, 25 miRNAs were upregulated, and 23 miRNAs were downregulated significantly (Fig. 2B). Gene filter analysis of total miRNAs pool shows that 99.28% of miRNA population did not show significant difference in the cytosol vs synaptosome compartments. Only, 0.38% population of miRNAs is upregulated, and 0.35% miRNA population is downregulated (Fig. 2C). The scattered plot shows the average log2 fold changes values of significantly deregulated miRNAs (SI Fig. 2A) and the volcano plot shows the p values (-log10) of significantly deregulated miRNAs (SI Fig. 2B). The top candidate miRNAs were selected for validation analysis.

**Figure 2.**
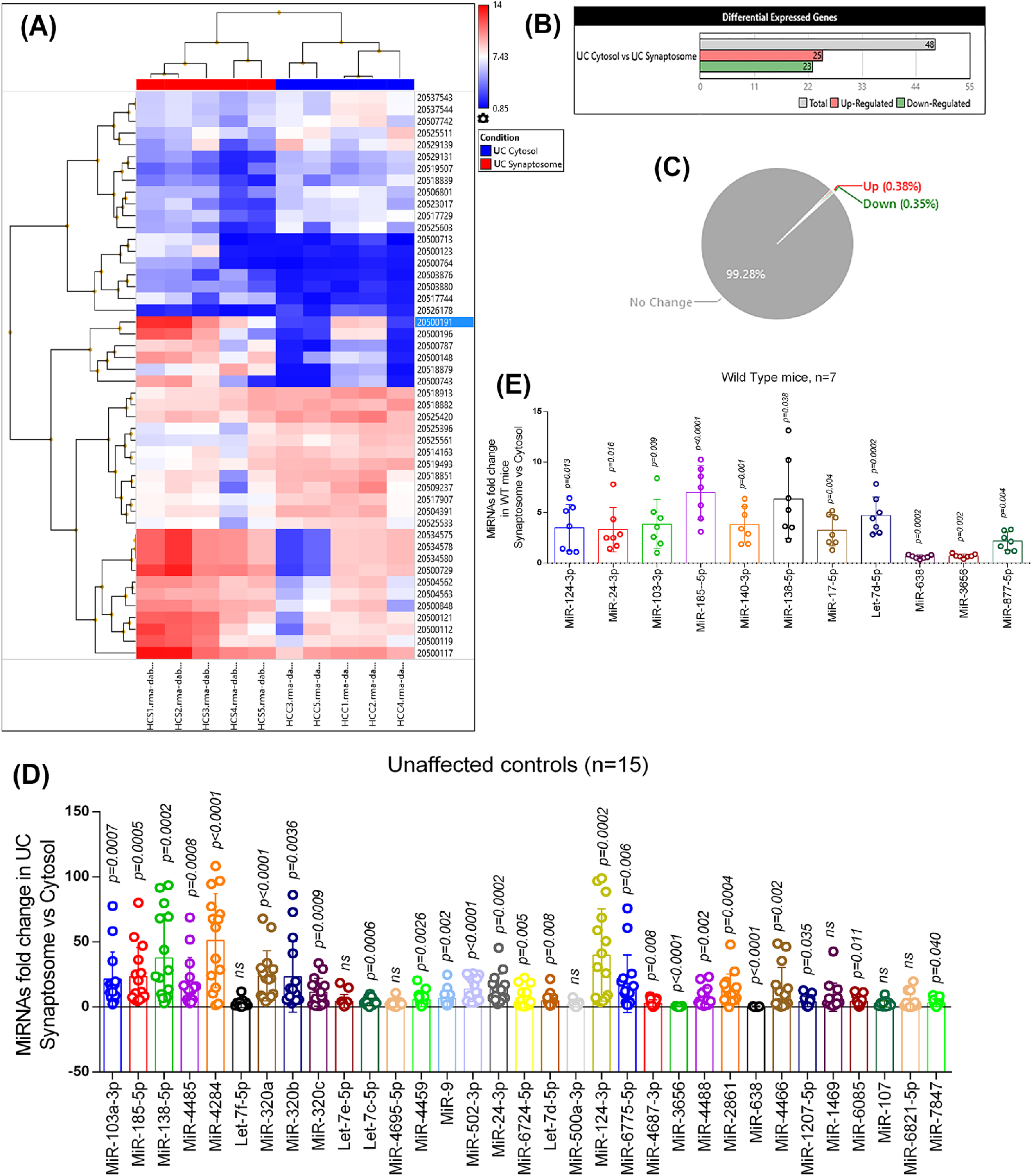
MiRNAs expression in synaptosome and cytosol in healthy state. **(A)** Hierarchical clustering and heat map of significantly deregulated miRNAs in the synaptosome and cytosol of unaffected controls. (Red color intensity showed the miRNAs upregulation and blue color intensity showed the miRNAs downregulation) **(B)** Total number of miRNAs deregulated in cytosol vs synaptosome in unaffected controls. (Gray scale bar- total number of miRNAs; Red scale bar- Up-regulated miRNAs; Green scale bar- Down-regulated miRNAs) **(C)** Pi diagram showed the total miRNAs pool distribution and percentage of miRNAs population changed in cytosol and synaptosome in unaffected controls. **(D)** qRT-PCR based validation analysis of significantly deregulated miRNAs in unaffected controls (n=15). MiRNAs expression was quantified in terms of fold changes in unaffected controls synaptosomes compared to cytosol. Each circle dot represents one sample. **(E)** Validation analysis of significantly deregulated mmu-miRNAs in WT mice (n=7). MiRNAs expression was quantified in synaptosome relative to cytosol. Each circle dot represents one animal.

### Validation analysis of synaptosomal and cytosolic miRNAs in healthy state

#### (i) UC postmortem brains

Validation analysis was performed on UC (n=15) postmortem brains to distinguish synaptosomal and cytosolic miRNAs in the normal state. Out of the 43 deregulated miRNAs, only 33 miRNAs were successfully amplified by qRT-PCR using specific primers. The 18 miRNAs showed similar expression trends as obtained by Affymetrix data analysis. The remaining miRNAs did not concur with Affymetrix data. Overall, 24 miRNAs were significantly upregulated in the synaptosomes relative to the cytosol and two miRNAs (miR-638 and miR-3656) were significantly downregulated in the synaptosomal fractions relative to the cytosolic fractions. Seven miRNAs did not show any significant changes (Fig. 2D).

#### (ii) WT mice brains

Further, we performed expression analysis of above classified synaptosomal and cytosolic miRNAs in WT mice (n=7). A total of 11 Mus musculus (mmu)-miRNAs were amplified, and out of them, nine were significantly upregulated and two were downregulated in WT mice synaptosome relative to cytosol (Fig. 2E). The 11 miRNAs showed similar expression pattern as observed by primary screening and UC postmortem brain validation. Based on these observation nine miRNAs were classified as synaptosomal miRNAs and two miRNAs as cytosolic miRNAs in the healthy state.

### MicroRNAs expression in AD synaptosomes vs AD cytosol

Next, we compared the microarray data for miRNAs expression changes in AD synaptosomal fractions vs AD cytosolic fractions. A total of 39 mature miRNAs were found to be deregulated in AD synaptosome vs AD cytosol comparison as shown in SI Table 5 and 28 hsa-miRNAs were highly expressed in the synaptosomes and low in the cytosol. The 11 out 39 miRNAs were highly expressed in the cytosol and showed reduced expression in the synaptosomes. The differential expression of these miRNAs in the AD synaptosomes and AD cytosol suggests their functional relevance in diseased state.

Fig. 3A shows hierarchical clustering and a heat map of significantly deregulated miRNAs with their ID numbers. The 11 miRNAs were upregulated in the cytosol and 28 miRNAs were downregulated in the cytosol significantly (Fig. 3B). Gene filter analysis of total miRNAs pool shows that 99.41% of miRNA population did not show significant difference in the cytosol vs synaptosome compartment, only, 0.59% of populations showed variable expression levels. The 0.17% of miRNAs are upregulated and 0.42% of miRNAs population is downregulated (Fig. 3C). The scattered plot shows the average log2 fold changes values of significantly deregulated miRNAs (SI Fig. 2C) and the volcano plot shows the p values (-log10) of significantly deregulated miRNAs in AD synaptosome vs AD cytosol (SI Fig. 2D). Based on the miRNA(s) expression pattern in unaffected controls and AD samples, 22 miRNAs (37.3%) were expressed only in UC samples and 21 miRNAs (35.6%) were expressed only in AD samples. However, 16 miRNAs (27.1%) were commonly expressed in both conditions (SI Fig. 3).

**Figure 3.**
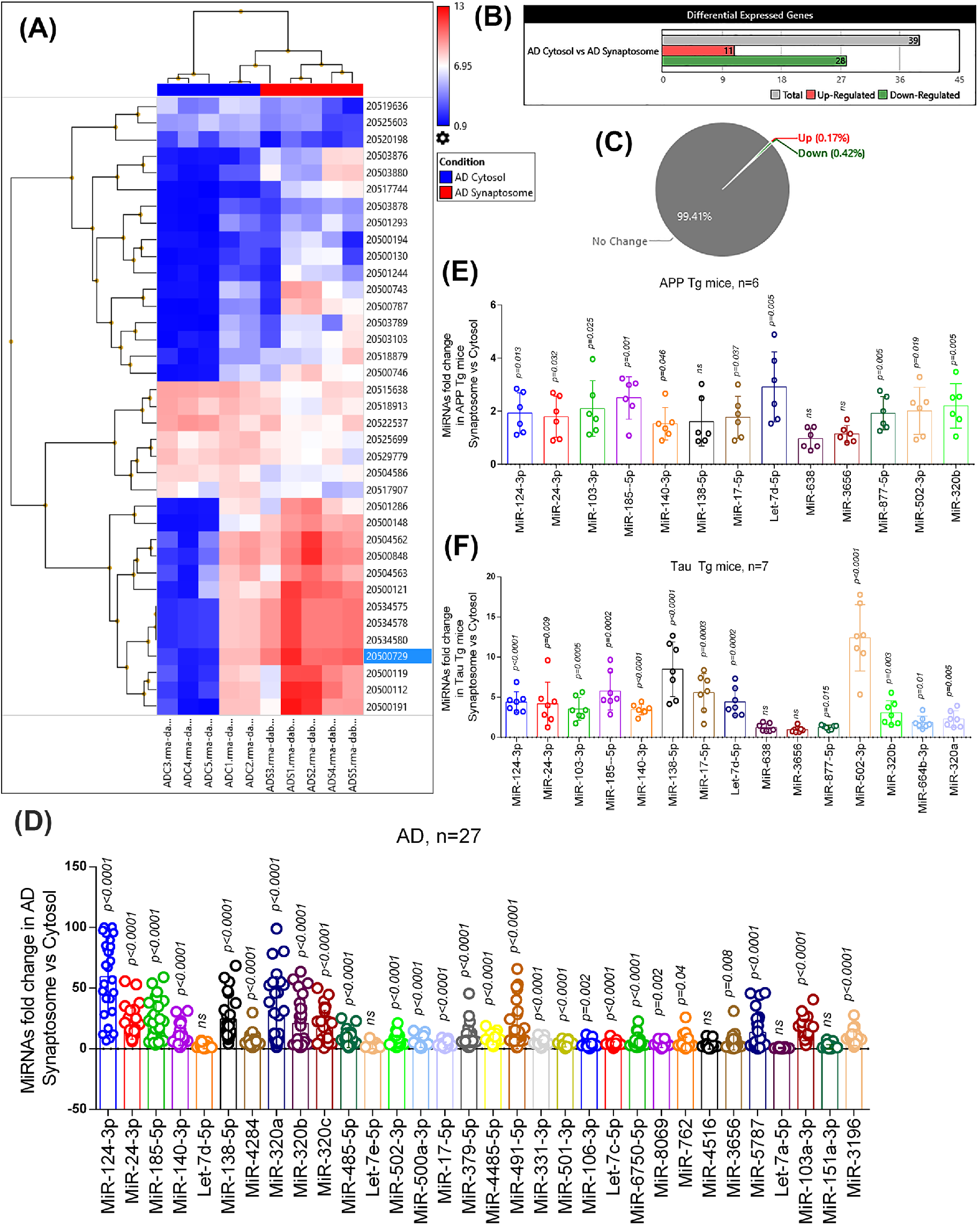
MiRNAs expression in synaptosome and cytosol in AD. **(A)** Hierarchical clustering and heat map of significantly deregulated miRNAs in cytosol and synaptosome in AD samples. (Red color intensity showed the miRNAs upregulation and blue color intensity showed the miRNAs downregulation) **(B)** Total numbers of miRNAs deregulated in cytosol and synaptosome in AD. (Gray scale bar- total number of miRNAs; Red scale bar- Up-regulated miRNAs; Green scale bar- Down-regulated miRNAs) **(C)** Pi diagram showed the total miRNAs pool distribution and percentage of miRNA populations changed in cytosol and synaptosome. **(D)** qRT-PCR based validation analysis of significantly deregulated miRNAs in AD samples (n=27). MiRNAs expression was quantified in terms of fold changes in AD synaptosome compared to AD cytosol. Each circle dot represents one sample. **(E)** Validation analysis of significantly deregulated mmu-miRNAs in APP-Tg (n=6) mice. MiRNAs expression was quantified in synaptosome relative to cytosol. Each circle dot represents one animal. **(F)** Validation analysis of significantly deregulated mmu-miRNAs in Tau-Tg (n=7) mice. MiRNAs expression was quantified in synaptosome relative to cytosol.

### Validation analysis of synaptosomal and cytosolic miRNAs in AD state

#### (i) AD postmortem brains

The top candidate miRNAs were selected for validation analysis. Validation analyses were performed on 27 AD postmortem brains to distinguish synaptosomal and cytosolic miRNAs in the diseased state. Out of the 39 deregulated miRNAs, 32 miRNAs were amplified by using specific primers. The 22 miRNAs showed the similar expression trend as obtained by Affymetrix data analysis. The remaining miRNAs either showed opposite trend to Affymetrix data or did not change significantly. Overall, 27 miRNAs were significantly upregulated in the synaptosomes relative to the cytosol and no miRNA showed any significantly downregulation. The five miRNAs did not show any significant changes in the synaptosomes relative to the cytosol (Fig. 3D).

#### (ii) APP-Tg mice

Next, we did synaptosomal and cytosolic miRNAs validation using APP-Tg mice (n=6). The 13 mmu-miRNAs showed similar expression pattern as observed by primary screening and AD postmortem brain validation. MiR-103-3p, miR-185-5p, miR-24-3p, miR-502-3p, miR-320b, let-7d-5p, miR-124-3p, miR-140-3p, miR-17-5p, and miR-877-5p showed significant upregulation in the synaptosomes, while miR-138-5p, miR-3656 and miR-638 did not show any significantly changes in their expression (Fig. 3E).

#### (iii) Tau-Tg mice

Further, we did synaptosomes and cytosolic miRNAs validation using Tau-Tg mice (n=7). The 13 mmu-miRNAs showed similar expression pattern as observed by primary screening and AD postmortem brain validation. MiR-103-3p, miR-185-5p, miR-24-3p, miR-502-3p, miR-320b, let-7d-5p, miR-124-3p, miR-140-3p, miR-17-5p, miR-877-5p, miR-320a and miR-664a-3p showed significant upregulation in the synaptosomes, while miR-138-5p, miR-3656 and miR-638 did not show any significantly changes in their expression (Fig. 3F).

Based on these observations, 11 miRNAs were classified as synaptosomal miRNAs and two miRNAs as cytosolic miRNAs in the AD state.

### MicroRNAs expression in AD cytosol vs UC cytosol

Next, we compared AD cytosolic vs UC cytosolic miRNAs. A total of 13 hsa-miRNAs were found to be significantly deregulated in the AD cytosol vs UC cytosol comparison SI Table 6. Interestingly, expression levels of all miRNAs were reduced in AD cytosol as mentioned in SI Table 6. SI Fig. 4A shows the hierarchical clustering and heat map of significantly deregulated miRNAs with their ID numbers. The 13 miRNAs were found to be downregulated significantly (SI Fig. 4B). Gene filter analysis of total miRNAs pool showed that 99.76% of miRNA population did not show significant difference in the cytosol vs synaptosome compartment. Only, 0.24% of miRNA populations showed variable expression levels. All 0.24% miRNA population is downregulated (SI Fig. 4C). The scattered plot showed the average log2 fold changes values of significantly deregulated miRNAs (SI Fig. 5A) and volcano plot showed the p values (-log10) of significantly deregulated miRNAs in AD cytosol vs AD cytosol (SI Fig. 5B). The top candidate miRNAs were selected for validation analysis.

### Validation analysis of cytosolic miRNAs in AD and unaffected control

#### (i) AD and UC postmortem brains

Validation analysis of cytosolic miRNAs were performed on 15 UC and 27 AD postmortem brain samples. The 13 miRNAs candidates were selected for validation analysis. Opposed to the Affymetrix data, nine miRNAs were significantly upregulated in AD cytosol relative to UC cytosol and three miRNAs did not show significant changes (SI Fig. 4D).

#### (ii) WT, APP-Tg and Tau-Tg mice

We also performed the validation of cytosolic miRNAs in APP-Tg and Tau-Tg mice relative to WT mice. Other than the 13 cytosolic mmu-miRNAs, we also checked the expression of other potential mmu-miRNAs: miR-17-5p, let-7d-5p, miR-185-5p, miR-103-3p, miR-138-5p, miR-877-5p, miR-24-3p, miR-502-3p, miR-140-3p, miR-124-3p and miR-3656. Most of the miRNAs were upregulated in the APP-Tg and Tau-Tg cytosol relative to WT cytosol. Only, miR-638 and miR-3656 were significantly down regulated in APP-Tg cytosol relative to WT (SI Fig. 6).

### MicroRNAs expression in AD synaptosomes vs UC synaptosomes

Lastly, we compared the microarray data for miRNAs expression changes in AD synaptosomes vs UC synaptosomes. A total of 11 miRNAs were found to be deregulated significantly in AD synaptosomes vs UC synaptosomes comparison as shown in (SI Table 7). Four hsa-miRNAs-miR-502-3p, miR-500a-3p, miR-877-5p and miR-664b-3p were highly expressed in AD synaptosomes relative to UC synaptosomes. The remaining seven hsa-miRNAs-miR-3196, miR-6511b-5p, miR-4508, miR-1237-5p, miR-5001-5p, miR-4492 and miR-4497 showed reduced expression in AD synaptosomes and were highly expressed in UC synaptosomes. The differential expression of these miRNAs in AD and UC synaptosomes suggests their importance in synapse function.

Fig. 4A showed the hierarchical clustering and heat map of significantly deregulated miRNAs with their ID numbers. The four miRNAs were upregulated, and seven miRNAs were downregulated significantly (Fig. 4B). Gene filter analysis of total miRNAs pool showed that 99.83% of the miRNA population did not show significant difference in the synaptosome compartments in AD vs UC. Only 0.17% miRNAs populations showed variable expression pattern. The 0.06% of miRNAs is upregulated and 0.11% of the miRNA population is downregulated (Fig. 4C). The scattered plot showed the average log2 fold changes values of significantly deregulated miRNAs (SI Fig. 5C) and the volcano plot showed the p values (-log10) of significantly deregulated miRNAs in AD synaptosomes vs UC synaptosomes (SI Fig. 5D). The top candidate miRNAs were selected for validation analysis.

**Figure 4.**
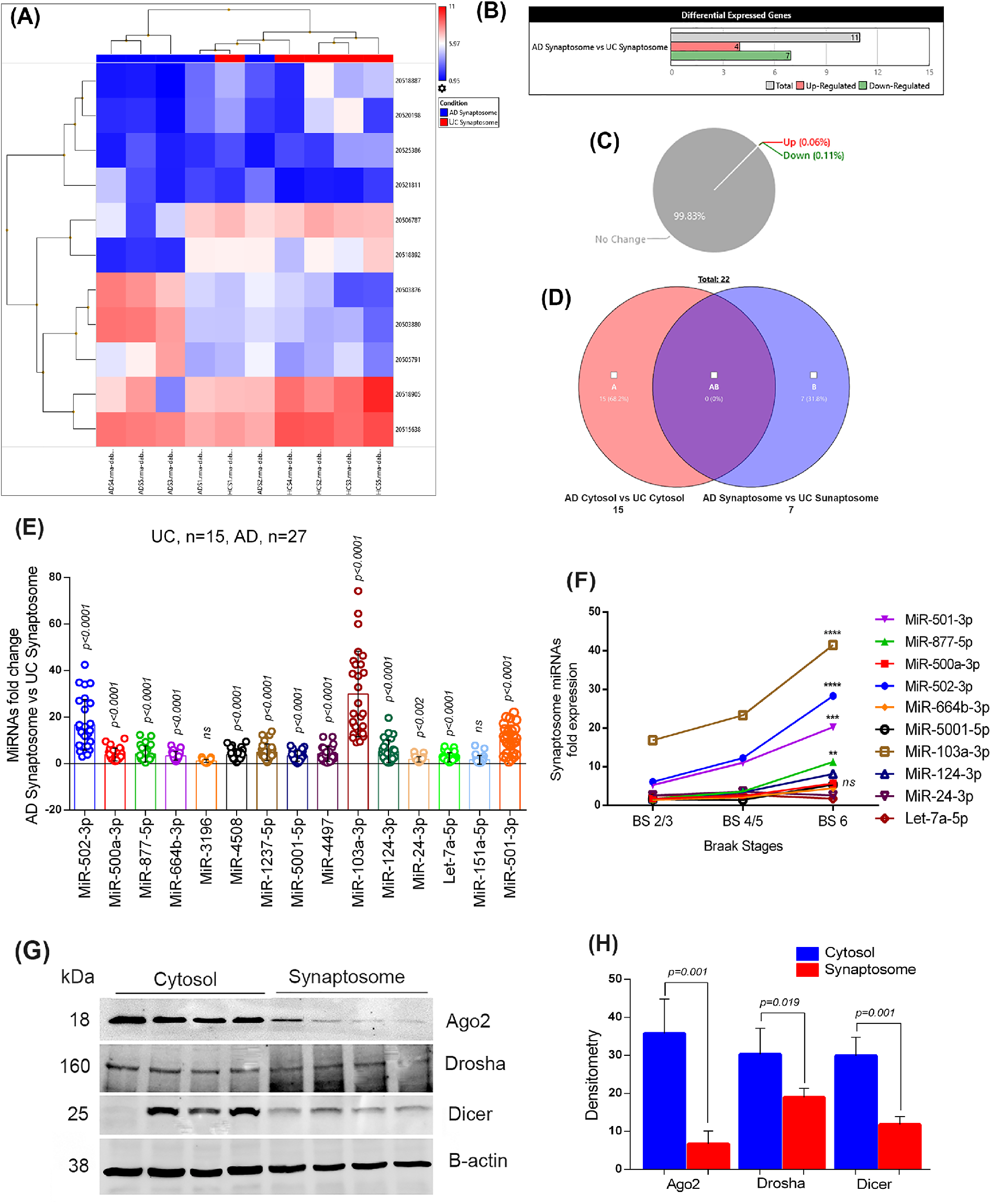
MiRNAs expression in synaptosome in AD and healthy state. **(A)** Hierarchical clustering and heat map of significantly deregulated miRNAs in synaptosome in AD and unaffected controls. (Red color intensity showed the miRNAs upregulation and blue color intensity showed the miRNAs downregulation) **(B)** Total numbers of miRNAs deregulated in AD synaptosome vs UC synaptosome. (Gray scale bar- total number of miRNAs; Red scale bar- Up-regulated miRNAs; Green scale bar- Down-regulated miRNAs) **(C)** Pi diagram showed the total miRNAs pool distribution and percentage of miRNAs population changed in AD synaptosome vs UC synaptosome. **(D)** Venn diagram showing the number of miRNAs that expressed only in cytosol and synaptosome in AD vs healthy state. **(E)** qRT-PCR based validation analysis of significantly deregulated miRNAs in AD (n=27) and UC (n=15) synaptosome. MiRNAs expression was quantified in terms of fold changes in AD synaptosome relative to UC synaptosome. Each circle dot represents one sample. **(F)** Multiple comparison analysis of synaptosomal miRNAs fold changes with Braak stages 2/3, Braak stages 4/5 and Braak stages 6 of AD samples. (**\*\***p<0.01, **\*\*\***p<0.001, **\*\*\*\***p<0.0001). **(G)** Immunoblotting analysis of miRNAs biogenesis proteins (Ago2, Drosha and Dicer) in cytosol and synaptosomal of UC samples (n=4). **(C)** Densitometry analysis of Ago2, Drosha and Dicer in cytosol relative to synaptosomes of UC samples.

Based on the miRNAs’ expression pattern in cytosol and synaptosomes in AD vs UC samples, 15 miRNAs (68.2%) were expressed only in the cytosol and seven miRNAs (31.8%) were expressed only in the synaptosomes. We did not see any miRNA that were commonly expressed in both conditions (Fig. 4D).

### Validation analysis of synaptosomal miRNAs

#### (i) AD and UC postmortem brains

Validation analysis were performed on 15 UC and 27 AD postmortem brains. We checked synaptosomal expression of deregulated 16 miRNAs. However, only 14 hsa-miRNAs were amplified, the 12 hsa-miRNAs (miR-502-3p, miR-500a-3p, miR-877-5p, miR-664b-3p, miR-4508, miR-1237-5p, miR-5001-5p, miR-4497, miR-103a-3p, miR-124-3p, miR-24-3p and let-7a-5p were significantly upregulated in the AD synaptosomes relative to UC synaptosomes, while two hsa-miRNAs (miR-3196 and miR-151-5p) did not show any significant changes (Fig. 4E).

#### (ii) WT, APP-Tg and Tau-Tg mice

We also performed the validation of above-mentioned miRNAs and other potential synaptosomal miRNAs in APP-Tg and Tau-Tg mice relative to WT mice. The 12 mmu-miRNAs, which were, amplified successfully included-miR-17-5p, let-7d-5p, miR-185-5p, miR-103-3p, miR-138-5p, miR-877-5p, miR-24-3p, miR-502-3p, miR-140-3p, miR-124-3p, miR-638 and miR-3656. In APP-Tg mice synaptosomes, seven miRNAs were significantly upregulated, four were significantly downregulated relative to WT synaptosomes and one miRNA showed no change (SI Fig. 6). In Tau-Tg synaptosomes, nine miRNAs were significantly upregulated, and three miRNAs were significantly downregulated relative to WT synaptosomes (SI Fig. 6).

Summarizing all validation analysis, 12 miRNAs expression was consistent in different comparisons and samples settings. The 10 miRNAs can be classified as synaptosomal miRNAs and two miRNAs as cytosolic miRNAs.

Next, we examined the synaptosomal miRNAs expression patterns with AD samples Braak stages. Multiple comparison analysis showed that expression of synaptosomal miRNAs were gradually increased with Braak stages. However, significant differences were found in miR-501-3p (p=0.001), miR-502-3p (p<0.0001), miR-877-5p (p=0.010) and miR-103a-3p (p<0.0001) fold changes at Braak stage 6 relative to Braak stage 2/3 (Fig. 4F). These results unveiled the strong connection of these miRNAs with AD progression.

Further, to determine the synaptosomal miRNAs synthesis at synapse, we checked the levels of key miRNA biogenesis proteins (Ago2, Drosha and Dicer) in cytosol and synaptosome fractions. In Fig. 4G, immunoblots showed the levels of miRNA biogenesis proteins in UC cytosol and synaptosomes. Densitometry analysis showed very high levels of all three proteins in cytosol relative to synaptosomes (Fig. 4H). The presence of miRNA biogenesis proteins in synaptosomes confirmed that miRNAs might be synthesize at synapse.

### In-silico Ingenuity® Pathway Analysis of cytosolic and synaptosomal miRNAs in AD and healthy state

The deregulated miRNAs under different conditions were run for IPA analysis. The first comparison was cytosolic vs synaptosomal miRNAs in the healthy state. The top deregulated miRNAs were involved in several diseases, molecular and cellular functions, physiological system development and functions (SI Table 8 and SI Table 9). However, we focused on the miRNA candidates which are involved in nervous system development and function in neurological diseases. Eleven miRNAs were identified which were significantly (P<0.05) involved in many neurological diseases and dementia including AD and MCI (SI Fig. 7A). Next, we analyzed the mRNA target and seed sequences of these miRNAs to understand the molecular mechanism of miRNAs involved in neurological function (SI Fig. 7B). The tumor suppressor gene (TP53) was the central gene that was targeted by many of these miRNAs. Other potential genes were BACE1, Smad2/3, Lypla1, Akt1 and SERBP1 pathway genes.

Similarly, we studied synaptosomal and cytosolic miRNAs function in AD cases. The top miRNA candidates were significantly (P<0.05) involved in several nervous system development, function and neurological diseases (SI Table 10). However, our interest was neurological disorders and dementia, where eight miRNAs were detected which were involved in several neurological disorders including AD (Fig. 5A). Further, miRNAs target predication analysis showed more than 20 genes that are targeted by these miRNAs (Fig. 5B). Next, we studied the biological roles of cytosolic miRNAs which were downregulated in AD compared to UC. The top five miRNAs were significantly involved in several diseases and molecular pathways (SI Table 11). MiRNAs and diseased pathways showed integration with Amyotrophic lateral sclerosis (SI Fig. 8A). Like other miRNAs, several genes were identified as potential target for these five miRNAs (SI Fig. 8B).

**Figure 5.**
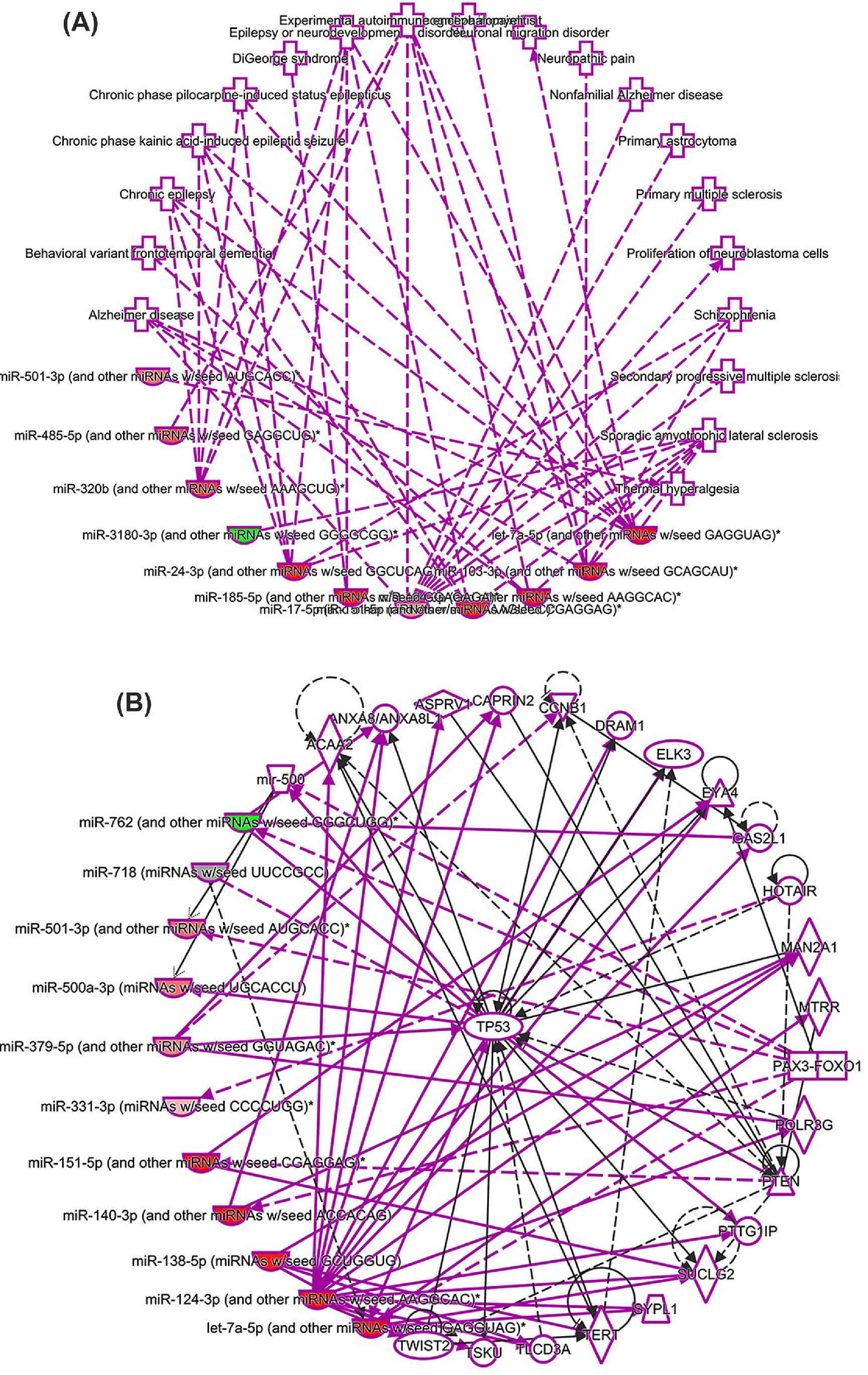
Ingenuity Pathway Analysis of cytosolic and synaptosomal miRNAs in AD. **(A)** In AD state-cytosolic and synaptosomal miRNAs expression network in various human diseases. Green nodes represent decreased expression and red nodes represent increased expression of miRNAs. **(B)** MiRNAs target and seed sequences network of cytosolic and synaptosomal miRNAs in the AD state.

Lastly, we studied the biological functions of synaptosomal miRNAs which were deregulated in AD vs UC. The miR-500 family (miR-501-3p, miR-500a-3p) and miR-877-5p were identified to be significantly (P<0.05) involved in several biological process and disorders (SI Table 12). MiRNA and disease interaction analysis showed a significant connection of miR-501-3p in GABAergic synapse function and other brain functions (Fig. 6A). The miRNAs target predication analysis showed more than 20 genes that are targeted by these miRNAs (Fig. 6B). The KRAS gene was identified as one of the potential common target of miR-501-3p, miR-502-3p and miR-877-5p (SI Fig. 9). Further, gene ontology enrichment analysis of miR-502-3p showed that it involved in several biological processes, cellular components and molecular functions. The most significant involvement was response to external stimuli (P=0.009) and nervous system development (P=0.044). The most significant cellular component was GABAergic synapse (P=0.028) and molecular function was calmodulin binding (P=0.020) (SI Fig. 10).

**Figure 6.**
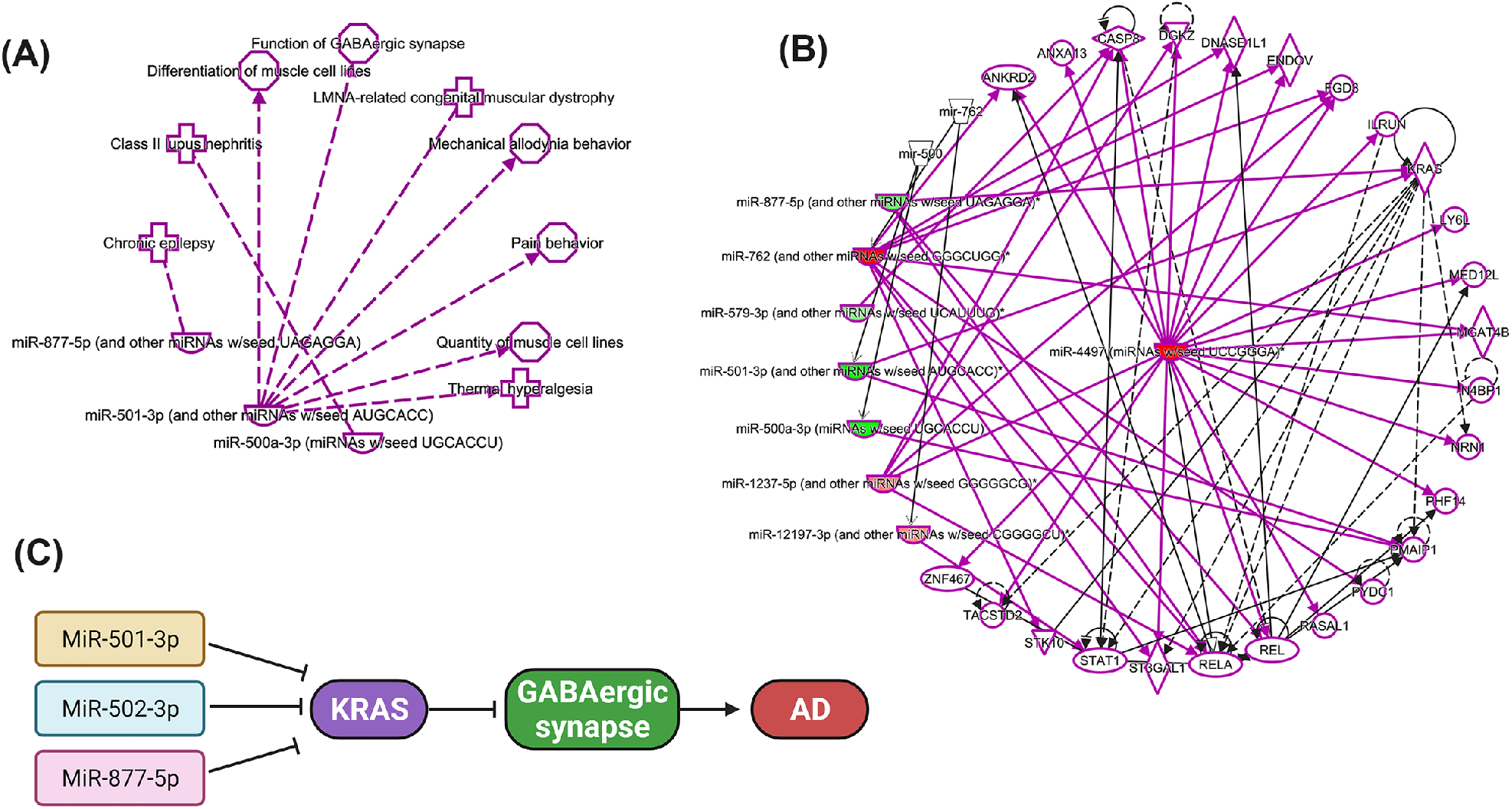
Ingenuity Pathway Analysis of synaptosomal miRNAs in AD. **(A) S**ynaptosomal miRNAs expression network in various human diseases. **(B)** MiRNAs target and seed sequences network of synaptosomal miRNAs in the AD and healthy state. Green nodes represent increased expression and red nodes represent decreased expression of miRNAs. **(C)** Possible molecular mechanism of GABAergic synapse regulation by miR-501-3p, miR-502-3p and miR-877-5p. KRAS gene is one of the top predicted target of miR-501-3p, miR-502-3p and miR-877-5p. Inhibition of KRAS expression by the overexpression of these miRNAs could inhibits the GABAergic synapse function in AD.

Overall, IPA and gene ontology enrichment analyses showed that synaptosomal miRNAs are altered in several neurological disorders and participate in numerous cellular and molecular pathways related to brain function.

## Discussion

Synaptosome based research in AD began since the discovery of the synaptosome by Hebb and Whittaker in 1958 (Hebb and Whittaker 1958). Although significant research has been done on synaptosomal function/dysfunction, we still know very little about physiological connections and pathological changes in AD, particularly the sequence of events that occur at synapse and the regulation of miRNAs in synaptosomes and how synaptosomal miRNAs are different from cytosolic miRNAs.

Using global synaptosomal and cytosolic miRNA analysis, in-silico analysis, transmission electron microscopy of healthy unaffected and AD postmortem brains and brain tissues from APP and Tau transgenic mice, in the current study we investigated a comprehensive synaptic and cytosolic miRNAs analysis. We also determined the possible molecular function of synaptic miRNAs in AD and brain aging.

It is well studied that miRNAs are present in different cell organelles and cellular components such as the nucleus, mitochondria, Golgi bodies, exosomes and apoptotic bodies. These differentially expressed miRNAs, can modulate the levels of localized proteins (Jie et al. 2021). Therefore, we hypothesized that synapse centered miRNAs are altered in AD. We also hypothesize that miRNAs in synaptosomes and cytosols are ‘differently expressed and localized’ in healthy (unaffected controls) and AD states. Therefore, for the first time, our study distinguished cytosolic and synaptosomal miRNAs and their alterations in healthy and AD states.

We examined cytosolic and synaptosomal miRNAs changes in both healthy and disease states. In primary screenings, some individual synaptosomal and cytosolic miRNAs were identified as those which were expressed in both healthy and disease states but with varying expression levels, in terms of fold change (≤ -2 and ≥ 2). We noted that fold change of similar synaptosomal miRNAs varied by >100-folds in AD relative to healthy state. Most of these synaptosomal miRNAs are studied in human diseases, but very limited information is available on the cytosolic miRNAs.

Interestingly, as shown by pie chart analysis, >99% of miRNAs population did not show significant changes in the synaptosome and cytosol. Only a small fraction (<1%) of miRNA pool showed significant changes among synaptosomes and cytosol populations. These findings confirmed that most of the miRNA populations are uniformly distributed in the neuron with an exception of some localized synapse miRNAs. These synaptosomal miRNAs are either synthesized locally at the synapse or may be transported from the soma to the synapse. As per our initial analysis, it seems that miRNA biogenesis machinery is present at the synapse, and it is possible that miRNAs processing occurs at the synapse. However, additional research is needed to confirm miRNA biogenesis at the synapse.

Validation analysis on the postmortem brains and brain tissues from AD mouse models amplified only limited numbers of miRNAs compared to primary Affymetrix screening. Our extensive and careful validation analysis of postmortem brains revealed several potential miRNAs that showed similar expression trends specified as synaptosomal or cytosolic miRNAs in both healthy and AD states. Further, extended validation analysis of APP-Tg and Tau-Tg mice shortlisted quite a few specific miRNAs. Overall, human and mouse data analyses revealed ten potential miRNAs designated as synaptosomal miRNAs shown in S Figure 6 are actively involved in several neural functions (Kang et al. 2017; Hara et al. 2017; Yang et al. 2018; Boscher et al. 2021; Estfanous et al. 2021; Liang et al. 2021).

Interesting data was obtained in the case of cytosolic miRNAs in AD vs healthy controls. The initial screening showed reduced expression of all cytosolic miRNAs in AD cytosol. This could be because of higher Aβ and p-tau concentration in the cytoplasm compared to the synapse and high toxicities may be responsible for altered expression of miRNAs. Our careful validation analysis using postmortem brains, WT mice, APP-Tg and Tau-Tg mice strongly unveiled miR-638 and miR-3656 as potential cytosolic miRNAs. Both miRNAs are unique in AD and need further investigation on cytosolic basis of AD progression.

The top synaptosomal miRNAs are miR-500a-3p, miR-501-3p, miR-502-3p and miR-877-5p. In addition, the most down regulated miRNA was miR-4499 as shown by the primary screening. MiR-500 cluster miRNAs were amplified in all validation settings; however, we did not see any significant expression of miR-4499 in the validation phase. The Gene Ontology Enrichment and IP analysis for the miR-500 cluster showed that miR-500 family is involved in key biological process, cellular function and molecular function.

The most significant biological process is response to external stimulus and the most significant cellular component is GABAergic synapse (SI Fig. 10). GABAergic synapse is a crucial inhibitory synapse that is dysfunctional in AD (Govindpani et al. 2017; Hollnagel et al. 2019; Xu et al. 2020; Jiménez-Balado and Eich 2021). Our results also confirmed reduced levels of GABRA1 in AD synaptosomes. Further, in-silico analysis showed that miR-502-3p could modulate the function of GABAergic synapse. Both Gene Ontology and IP analysis confirmed the strong links of miR-501-3p and miRF-502-3p in GABAergic synapse pathways. It could be mediated via modulation of the KRAS gene by these miRNAs (Fig. 6C). Further, miR-501-3p, miR-502-3p and miR-877-5p expression was significantly increased with Braak stages of AD postmortem brains again confirming the strong connection of these miRNAs with AD. Therefore, more research is warranted to study the roles of miR-501-3p, miR-502-3p and miR-877-5p in the regulation of excitatory and inhibitory synapse function in relation to AD.

In summary, our study identified the synaptosomal miRNAs that are deregulated in AD. Our comprehensive analysis identified the three most promising synaptosomal miRNAs- miR- 501-3p, miR-502-3p and miR-877-5p that could modulate the function of excitatory and inhibitory synapses in AD. Our ongoing research investigating the underlying molecular mechanism of miR-501-3p and miR-502-3p in synaptic activity and GABAergic synapse function in relation to Aβ and p-tau induced toxicities.

## Materials and methods

### Postmortem brain samples

Postmortem brains from AD patients and unaffected controls were obtained from NIH NeuroBioBanks- (1) Human Brain and Spinal Fluid Resource Center, 11301 Wilshire Blvd (127A), Los Angeles, CA. (2) Brain Endowment Bank, University of Miami, Millar School of Medicine, 1951, NW 7th Avenue Suite 240, Miami, FL. (3) Mount Sinai NIH Brain and Tissue Repository, 130 West Kingsbridge Road Bronx, NY (Kumar and Reddy 2018). Brain tissues were dissected from the Brodmann’s Area 10 of the frontal cortices from AD patients (n = 27) and age and sex matched unaffected controls (n = 15). Demographic and clinical details of study specimens are provided in SI Table 1.

### Synaptosomes extraction

Synaptosomes were extracted using Syn-PER Reagent as per manufacturer instructions with some modifications (Thermo Scientific, USA) (Zolochevska and Taglialatela 2020; Yoshino et al. 2021; Franklin and Taglialatela 2016). Briefly, 50 mg of brain tissue was used from each sample for synaptosome extraction in 1 ml of Syn-PER Reagent. Tissues were homogenized slowly by Dounce glass homogenization on ice with ∼10 slow strokes. The resulting tissue homogenates were transferred to a centrifuge tube. Samples were centrifuged at 1400 × g for 10 minutes at 4°C to remove the leftover tissue debris. After centrifugation, the supernatant was transferred to a new tube. Again, supernatant (homogenate) was centrifuged at high speed 15,000 × g for 20 minutes at 4°C. The supernatant was removed as cytosolic fraction and synaptosomes recovered in the pellet form. Both the cytosolic fraction and synaptosome pellet were processed for RNA and protein extraction. The synaptosome pellet was also processed for transmission electron microscopic (TEM) analysis.

### Synaptosomes Characterization

Synaptosome preparations (purity and integrity) were characterized by TEM analysis of synapse assembly, immunoblotting of synaptic proteins-synapse associate protein 25 (SNAP25), PSD95 and synaptophysin, and qRT-PCR analysis of similar synaptic genes (Biesemann et al. 2014; Postupna et al. 2014).

### Transmission Electron Microscopy of synaptosomes

Freshly isolated synaptosomes were processed for TEM analysis. Briefly, the pellet was fixed in a solution of 0.1M cacodylate buffer, 1.5% paraformaldehyde and 2.5% glutaraldehyde and then post-fixed with 1% osmium tetroxide and embedded in LX-112 resin. Ultrathin sections were cut, stained with uranyl acetate and lead citrate, and examined with the Hitachi H-7650 /Transmission Electron Microscope at 60 kV located at the College of Arts and Sciences Microscopy, Texas Tech University. Low-magnification imaging was followed by high-magnification imaging. Representative images were acquired and recorded with an AMT digital camera (Kumar et al. 2019).

### Immunoblotting analysis

We performed immunoblot analysis for the synaptic/cytosolic proteins, brain cells and miRNAs biogenesis proteins. Details of the proteins and antibody dilutions are given in SI Table 2. The 40 µg of protein lysates were resolved on a 4-12% Nu-PAGE gel (Invitrogen). The resolved proteins were transferred to nylon membranes (Novax Inc., San Diego, CA, USA) and then incubated for 1 h at room temperature with a blocking buffer (5% dry milk dissolved in a TBST buffer). The nylon membranes were incubated overnight with the primary antibodies. The membranes were washed with a TBST buffer 3 times at 10-min intervals and then incubated for 2 h with an appropriate secondary antibody, sheep anti-mouse HRP 1:10,000, followed by three additional washes at 10-min intervals. Proteins were detected with chemiluminescence reagents (Pierce Biotechnology, Rockford, IL, USA), and the bands from the immunoblots were visualized (Kumar et al. 2019).

### Quantitative Real-Time PCR analysis

Quantification of mRNA levels of synaptic genes was carried out with real-time qRT-PCR using methods described in Kumar et al. 2019 [54]. The oligonucleotide primers were designed with primer express software (Applied Biosystems) for SNAP25, synaptophysin, PSD95, elF1a and PCNA. The primer sequences and amplicon sizes are listed in SI Table 3. SYBR-Green chemistry-based quantitative real-time qRT-PCR was used to measure mRNA expression of these genes using β-actin as housekeeping genes, as described previously (Kumar et al. 2019).

### Affymetrix miRNA microarray analysis

Initially, we used five AD postmortem and five unaffected control (UC) postmortem brains for Affymetrix microarray analysis. The demographic and clinical details of samples used for Affymetrix analysis are given in Table 1. Total RNA was extracted from the synaptosomal and cytosolic fractions from both AD and unaffected control samples using the TriZol reagent with some modifications. Total we had 20 samples for miRNA analysis-AD synaptosome (n=5), UC synaptosome (n=5), AD cytosol (n=5) and UC cytosol (n=5). Detailed miRNAs screening of the synaptosome and cytosolic miRNAs were conducted at the University of Texas Southwestern Medical Center, Genomics and Microarray Core Facility, Dallas. The miRNA expression profiles were generated with Affymetrix GeneChip miRNA array v. 4.0 (Supplementary information).

**Table 1.** Demographic and clinical details of postmortem brains used for Affymetrix microarray analysis.

| S. No. | Barcode | Age | Sex | Disease status | Race | Brain region | CD R | Braak Score | PMI (hrs) | Cause of Death |
| --- | --- | --- | --- | --- | --- | --- | --- | --- | --- | --- |
| 1 | 77423 | 79 | F | AD | W | BM-10 | 3 | 6 | 6.50 | Coronary Artery disease |
| 2 | 77424 | 69 | M | AD | W | BM-10 | 3 | 6 | 5.42 | SEPSIS |
| 3 | 77425 | 75 | M | AD | W | BM-10 | 2 | 6 | 8.00 | Respiratory failure |
| 4 | 77426 | 94 | F | AD | W | BM-10 | 5 | 6 | 4.33 | Acute Myocardial Infraction |
| 5 | 77427 | 82 | M | AD | W | BM-10 | 5 | 6 | 20.67 | Cardiorespiratory arrest |
| 6 | 77428 | 65 | M | Unaffected control | H | BM-10 | 0 | 0 | 3.83 | Renal failure |
| 7 | 77431 | 103 | F | Unaffected control | W | BM-10 | 0 | 1 | 3.83 | Lymphadenopathy |
| 8 | 77433 | 75 | M | Unaffected control | B | BM-10 | 0 | 1 | 5.00 | Myocardial infraction |
| 9 | 77436 | 93 | M | Unaffected control | W | BM-10 | 0.5 | 0 | 4.17 | Acute Myocardial Infraction |
| 10 | 77437 | 84 | F | Unaffected control | W | BM-10 | 0 | 1 | 5.48 | Arteriosclerotic Heart Disease |

### Microarray data analysis

Data was analyzed using four comparisons- 1) AD synaptosome vs AD cytosol, 2) unaffected control (UC) synaptosome vs UC cytosol, 3) AD cytosol vs UC cytosol, and 4) AD synaptosome vs UC synaptosome. Microarray data for miRNAs expression changes in synaptosomal vs cytosol fractions were analyzed using two main criteria’s- Gene-level fold change < -2 or > 2 and Gene-level P-value <0.05. A probe set (Gene/Exon) is considered expressed if ≥ 50% samples have detectable above background (DABG) values below DABG Threshold < 0.05.

### Validation of deregulated miRNAs using postmortem brains

The deregulated miRNAs obtained from Affymetrix analysis were further tested and validated on large number of AD postmortem brains (n=27) and unaffected controls (n=15). Validation of miRNAs were performed for four comparisons- 1) AD synaptosome vs AD cytosol, 2) UC synaptosome vs UC cytosol, 3) AD cytosol vs UC cytosol, and 4) AD synaptosome vs UC synaptosome. MiRNAs levels were quantified by using miRNAs qRT-PCR, which involved three steps (i) miRNAs polyadenylation, (ii) cDNA synthesis and (iii) qRT-PCR as described previously (Kumar et al. 2014; Kumar et al. 2017; Kumar and Reddy 2021). Primers for desired miRNAs were synthesized commercially (Integrated DNA Technologies Inc., IA, USA) (SI Table 3). To normalize the miRNA expression, U6 snRNA and sno-202 were used as internal controls. The reaction mixture of each sample was prepared in triplicates. The reaction was set in the 7900HT Fast Real Time PCR System (Applied Biosystems, USA). qRT-PCR was performed in triplicate, and the data were expressed as the mean ±SD.

### Validation of differentially expressed miRNAs using AD mouse models

The deregulated miRNAs obtained from Affymetrix analysis were further validated using brain tissues from 12-month-old APP Transgenic (Tg2576) (n=6), Tau transgenic (P301L) (n=7) and age and sex matched wild type (WT) (n=7) mice. The deregulated miRNAs were conserved in both human and mice. The APP-Tg, Tau-Tg and WT mice were obtained from Jackson Laboratories and the colonies were maintained in our lab. Mice were euthanized to extract brain tissues. The brains were dissected, and the cerebral cortex was used for cytosol and synaptosome miRNA extraction. Validation of miRNAs were performed for four comparisons- 1) AD mice synaptosome vs cytosol, 2) WT mice synaptosome vs cytosol, 3) AD mice cytosol vs WT mice cytosol, and 4) AD mice synaptosome vs WT mice synaptosome. MiRNAs levels in APP and Tau mice relative to WT mice were quantified by using miRNAs qRT-PCR.

### In-silico analysis for potential miRNAs

The QIAGEN’s Ingenuity® Pathway Analysis (IPA®, QIAGEN Inc., https://www.qiagenbioinformatics.com/products/ingenuity-pathway-analysis) program was used to analyze the synaptosomal and cytosolic miRNAs target genes with FDR p-values <0.05 and with p-value <0.05. The IPA was used to gain insight into the overall biological changes caused by the expression, target gene prediction for synaptosomal and cytosolic miRNAs with AD and unaffected controls and gene Integrated Analysis. Each gene was related to various functions, pathways, and diseases as analyzed using Ingenuity knowledge base platform. The miRNA target genes (predicted and validated) were identified using various on-line miRNA algorithms (diana-microt, microrna.org, mirdb, rna22-has, targetminer, and targetscan-vert) (Kumar et al. 2017; Vijayan et al. 2018).

### Statistical considerations

Statistical parameters were calculated using Prism software, v6 (La Jolla, CA, USA). Results are reported as mean ± SD. The results were analyzed by two-tailed Student’s t-test to evaluate miRNAs expression in two groups of samples- 1) AD synaptosome vs AD cytosol, 2) UC synaptosome vs UC cytosol, 3) AD cytosol vs UC cytosol, and 4) AD synaptosome vs UC synaptosome. One-way comparative analysis of variance was used for analyzing WT, APP-Tg and Tau-Tg mice synaptosome vs cytosolic miRNAs data. Significant differences in three group of samples were calculated by Bonferroni’s multiple comparison tests. The correlation of miRNAs fold changes with Braak stages were analyzed by Tukey’s multiple comparisons test. P <0.05 was considered statistically significant.

## Acknowledgements

The authors would like to thank NIH for funding various projects - R01AG042178, R01AG47812, R01NS105473, AG060767, AG069333, and AG066347 (P.H.R), P30AG AG072973 (R.H.S.), AG051086 (D.K.L.), and K99AG065645 (S.K.).

## References

1. 2021 Alzheimer’s disease facts and figures. 2021. Alzheimers Dement 17:327–406.

2. Forner S, Baglietto-Vargas D, Martini AC, Trujillo-Estrada L, LaFerla FM. 2017. Synaptic Impairment in Alzheimer’s Disease: A Dysregulated Symphony. Trends Neurosci 40:347–357.

3. Marsh J, Alifragis P. 2018. Synaptic dysfunction in Alzheimer’s disease: the effects of amyloid beta on synaptic vesicle dynamics as a novel target for therapeutic intervention. Neural Regen Res 13:616–623.

4. Kashyap G, Bapat D, Das D, Gowaikar R, Amritkar RE, Rangarajan G, Ravindranath V, Ambika G. 2019. Synapse loss and progress of Alzheimer’s disease -A network model. Sci Rep 9:6555.

5. Selkoe DJ. 2002. Alzheimer’s disease is a synaptic failure. Science 298:789–791.

6. Chen Y, Fu AKY, Ip NY. 2019. Synaptic dysfunction in Alzheimer’s disease: Mechanisms and therapeutic strategies. Pharmacol Ther 195:186–198.

7. Ahmad F, Liu P. 2020. Synaptosome as a tool in Alzheimer’s disease research. Brain Res 1746:147009.

8. Colom-Cadena M, Spires-Jones T, Zetterberg H, Blennow K, Caggiano A, Dekosky S, Fillit H, Harrison J, Catalano SM et al. 2020. The clinical promise of biomarkers of synapse damage or loss in Alzheimer’s disease. Alzheimers Res Ther 12:21.

9. Kumar S, Reddy PH. 2020. The role of synaptic microRNAs in Alzheimer’s disease. Biochim Biophys Acta Mol Basis Dis. 1866:165937.

10. Reddy PH, Tripathi R, Troung Q, Tirumala K, Reddy TP, Anekonda V, Shirendeb UP, Calkins M, Reddy AP, Manczak M et al. 2012. Abnormal mitochondrial dynamics and synaptic degeneration as early events in Alzheimer’s disease: implications to mitochondria-targeted antioxidant therapeutics. Biochim Biophys Acta 1822:639–49.

11. Spires-Jones TL, Hyman BT. 2014. The intersection of amyloid beta and tau at synapses in Alzheimer’s disease. Neuron 82:756–71.

12. Jackson J, Jambrina E, Li J, Marston H, Menzies F, Phillips K, Gilmour G. 2019. Targeting the Synapse in Alzheimer’s Disease. Front Neurosci 13:735.

13. Calkins MJ, Manczak M, Mao P, Shirendeb U, Reddy PH. 2011. Impaired mitochondrial biogenesis, defective axonal transport of mitochondria, abnormal mitochondrial dynamics and synaptic degeneration in a mouse model of Alzheimer’s disease. Hum Mol Genet 20:4515–29.

14. Calkins MJ, Manczak M, Reddy PH. 2012. Mitochondria-Targeted Antioxidant SS31 Prevents Amyloid Beta-Induced Mitochondrial Abnormalities and Synaptic Degeneration in Alzheimer’s Disease. Pharmaceuticals (Basel) 5:1103–19.

15. Swerdlow RH. 2020. The mitochondrial hypothesis: Dysfunction, bioenergetic defects, and the metabolic link to Alzheimer’s disease. Int Rev Neurobiol 154:207–33.

16. Weidling IW, Swerdlow RH. 2020. Mitochondria in Alzheimer’s disease and their potential role in Alzheimer’s proteostasis. Exp Neurol 330:113321.

17. Kodavati M, Wang H, Hegde ML. 2020. Altered Mitochondrial Dynamics in Motor Neuron Disease: An Emerging Perspective. Cells 9:1065.

18. Ammal Kaidery N, Ahuja M, Sharma SM, Thomas B. 2021. An Emerging Role of miRNAs in Neurodegenerative Diseases: Mechanisms and Perspectives on miR146a. Antioxid Redox Signal 35:580–594.

19. John A, Reddy PH. 2021. Synaptic basis of Alzheimer’s disease: Focus on synaptic amyloid beta, P-tau and mitochondria. Ageing Res Rev 65:101208.

20. O’Brien J, Hayder H, Zayed Y, Peng C. 2018. Overview of MicroRNA Biogenesis, Mechanisms of Actions, and Circulation. Front Endocrinol (Lausanne) 9:402.

21. Lugli G, Torvik VI, Larson J, Smalheiser NR. 2008. Expression of microRNAs and their precursors in synaptic fractions of adult mouse forebrain. J Neurochem. 106:650–61.

22. Xu J, Chen Q, Zen K, Zhang C, Zhang Q. 2013. Synaptosomes secrete and uptake functionally active microRNAs via exocytosis and endocytosis pathways. J Neurochem. 124:15–25.

23. Li H, Wu C, Aramayo R, Sachs MS, Harlow ML. 2015. Synaptic vesicles contain small ribonucleic acids (sRNAs) including transfer RNA fragments (trfRNA) and microRNAs (miRNA). Sci Rep 5:14918.

24. Boese AS, Saba R, Campbell K, Majer A, Medina S, Burton L, Booth, TF, Chong P, Westmacott G, Dutta S. et al. 2016. MicroRNA abundance is altered in synaptoneurosomes during prion disease. Mol Cell Neurosci 71:13–24.

25. Rylett RJ, Ball MJ, Colhoun EH. 1983. Evidence for high affinity choline transport in synaptosomes prepared from hippocampus and neocortex of patients with Alzheimer’s disease. Brain Res 289:169–175.

26. Rajmohan R, Reddy PH. 2017. Amyloid-Beta and Phosphorylated Tau Accumulations Cause Abnormalities at Synapses of Alzheimer’s disease Neurons. J Alzheimers Dis 57:975–99.

27. Lauterborn JC, Scaduto P, Cox CD, Schulmann A, Lynch G, Gall CM, Keene D, Limon A. 2021. Increased excitatory to inhibitory synaptic ratio in parietal cortex samples from individuals with Alzheimer’s disease. Nat Commun 12:2603.

28. Govindpani K, Calvo-Flores Guzmán B, Vinnakota C, Waldvogel HJ, Faull RL, Kwakowsky A. 2017. Towards a Better Understanding of GABAergic Remodeling in Alzheimer’s Disease. Int J Mol Sci 18:1813.

29. Hollnagel JO, Elzoheiry S, Gorgas K, Kins S, Beretta CA, Kirsh J, Kuhse J, Kann O, Kiss E. 2019. Early alterations in hippocampal perisomatic GABAergic synapses and network oscillations in a mouse model of Alzheimer’s disease amyloidosis. PLoS One 14:e0209228.

30. Xu Y, Zhao M, Han Y, Zhang H. 2020. GABAergic Inhibitory Interneuron Deficits in Alzheimer’s Disease: Implications for Treatment. Front Neurosci 14:660.

31. Jiménez-Balado J, Eich TS. 2021. GABAergic dysfunction, neural network hyperactivity and memory impairments in human aging and Alzheimer’s disease. Semin Cell Dev Biol 116:146–59.

32. Jhou JF, Tai HC. 2017. The Study of Postmortem Human Synaptosomes for Understanding Alzheimer’s Disease and Other Neurological Disorders: A Review. Neurol Ther 6:57–68.

33. Schratt G. 2009. microRNAs at the synapse. Nat Rev Neurosci. 10:842–849

34. Siegel G, Saba R, Schratt G. 2011. microRNAs in neurons: manifold regulatory roles at the synapse. Curr Opin Genet Dev 21:491–7.

35. Wingo TS, Yang J, Fan W, Min Canon S, Gerasimov ES, Lori A, Logsdon B, Yao B, Seyfried NT, Wingo AP et al. 2020. Brain microRNAs associated with late-life depressive symptoms are also associated with cognitive trajectory and dementia. NPJ Genom Med 5:6.

36. Reddy PH, Beal MF. 2008. Amyloid beta, mitochondrial dysfunction and synaptic damage: implications for cognitive decline in aging and Alzheimer’s disease. Trends Mol Med 14:45–53.

37. Smalheiser NR. 2014. The RNA-centred view of the synapse: non-coding RNAs and synaptic plasticity. Philos Trans R Soc Lond B Biol Sci 369:20130504.

38. Ye Y, Xu H, Su X, He X. 2016. Role of MicroRNA in Governing Synaptic Plasticity. Neural Plast 2016:4959523.

39. John A, Kubosumi A, Reddy PH. 2020. Mitochondrial MicroRNAs in Aging and Neurodegenerative Diseases. Cells 9:1345.

40. Gowda P, Reddy PH, Kumar S. 2021. Deregulated mitochondrial microRNAs in Alzheimer’s disease: Focus on synapse and mitochondria. Aging Research Review 73: 101529.

41. Lahiri DK, Maloney B. 2010. Beyond the signaling effect role of amyloid-ß42 on the processing of APP, and its clinical implications. Exp Neurol 225:51–54.

42. Long JM, Lahiri DK. 2011. MicroRNA-101 downregulates Alzheimer’s amyloid-β precursor protein levels in human cell cultures and is differentially expressed. Biochem Biophys Res Commun 404:88995.

43. Long JM, Maloney B, Rogers JT, Lahiri DK. 2019. Novel upregulation of amyloid-β precursor protein (APP) by microRNA-346 via targeting of APP mRNA 5′-untranslated region: implications in Alzheimer’s disease. Mol Psychiatry 3:345–63.

44. Chopra N, Wang R, Maloney B, Nho K, Beck JS, Pourshafie N, Niculescu A, Saykin AJ, Rinaldi C, Lahiri DK et al. 2020. MicroRNA-298 reduces levels of human amyloid-β precursor protein (APP), β-site APP-converting enzyme 1 (BACE1) and specific tau protein moieties. Mol Psychiatry.

45. Lukiw WJ. 2020. microRNA-146a signaling in Alzheimer’s disease (AD) and prion disease (PrD). Front Neurol 25:11:462.

46. Zhao Y, Jaber VR, LeBeauf A, Sharfman NM, Lukiw WJ. 2019. microRNA-34a (miRNA-34a) mediated down-regulation of the post-synaptic cytoskeletal element SHANK3 in sporadic Alzheimer’s disease (AD). Front Neurol 10:28.

47. Kumar S, Reddy PH. 2016. Are circulating microRNAs peripheral biomarkers for Alzheimer’s disease? Biochim Biophys Acta 1862:1617–27

48. Zolochevska O, Taglialatela G. 2020. Selected microRNAs Increase Synaptic Resilience to the Damaging Binding of the Alzheimer’s Disease Amyloid Beta Oligomers. Mol Neurobiol. 57:2232–2243.

49. Yoshino Y, Roy B, Dwivedi Y. 2021. Differential and unique patterns of synaptic miRNA expression in dorsolateral prefrontal cortex of depressed subjects. Neuropsychopharmacology 46:900–10.

50. Kumar S, Reddy PH. 2018. MicroRNA-455-3p as a Potential Biomarker for Alzheimer’s Disease: An Update. Front Aging Neurosci 10:41.

51. Franklin W, Taglialatela G. 2016. A method to determine insulin responsiveness in synaptosomes isolated from frozen brain tissue. J Neurosci Methods 261:128–34.

52. Biesemann C, Grønborg M, Luquet E, Wichert SP, Bernard V, Bungers SR, Cooper B, Varoqueaux F, Li L, Herzog E et al. 2014. Proteomic screening of glutamatergic mouse brain synaptosomes isolated by fluorescence activated sorting. EMBO J 33:157–70.

53. Postupna NO, Keene CD, Latimer C, Sherfield EE, Van Gelder RD, Ojemann JG, Montine TJ, Darvas M. 2014. Flow cytometry analysis of synaptosomes from post-mortem human brain reveals changes specific to Lewy body and Alzheimer’s disease. Lab Invest 94:1161–72.

54. Kumar S, Reddy AP, Yin X, Reddy PH. 2019. Novel MicroRNA-455-3p and its protective effects against abnormal APP processing and amyloid beta toxicity in Alzheimer’s disease. Biochim Biophys Acta Mol Basis Dis 1865:2428–40.

55. Kumar S, Chawla YK, Ghosh S, Chakraborti A. 2014. Severity of hepatitis C virus (genotype-3) infection positively correlates with circulating microRNA-122 in patients sera. Dis Markers 2014:435476.

56. Kumar S, Vijayan M, Reddy PH. 2017. MicroRNA-455-3p as a potential peripheral biomarker for Alzheimer’s disease. Hum Mol Genet 26:3808–22.

57. Kumar S, Reddy PH. 2021. Elevated levels of MicroRNA-455-3p in the cerebrospinal fluid of Alzheimer’s patients: A potential biomarker for Alzheimer’s disease. Biochim Biophys Acta Mol Basis Dis 1867:166052.

58. Vijayan M, Kumar S, Yin X, Zafer D, Chanana V, Cengiz P, Reddy PH. 2018. Identification of novel circulatory microRNA signatures linked to patients with ischemic stroke. Hum Mol Genet 27(13):2318–29.

59. Scherma M, Qvist JS, Asok A, Huang SC, Masia P, Deidda M, Wei YB, Soni RK, Fratta W, Melas PA et al. 2020. Cannabinoid exposure in rat adolescence reprograms the initial behavioral, molecular, and epigenetic response to cocaine. Proc Natl Acad Sci USA 117:9991–10002

60. Hebb CO, Whittaker VP. 1958. Intracellular distributions of acetylcholine and choline acetylase. J Physiol 142:187–96.

61. Jie M, Feng T, Huang W, Zhang M, Feng Y, Jiang H, Wen Z. 2021. Subcellular Localization of miRNAs and Implications in Cellular Homeostasis. Genes (Basel) 12:856.

62. Kang Q, Xiang Y, Li D, Liang J, Zhang X, Zhou F, Qiao M, Nie Y, He Y, Li Y. 2017. MiR-124-3p attenuates hyperphosphorylation of Tau protein-induced apoptosis via caveolin-1-PI3K/Akt/GSK3β pathway in N2a/APP695swe cells. Oncotarget 8:24314–26.

63. Hara N, Kikuchi M, Miyashita A, Hatsuta H, Saito Y, Kuwano R, et al. 2017. Serum microRNA miR-501-3p as a potential biomarker related to the progression of Alzheimer’s disease. Acta Neuropathol Commun 5:10.

64. Yang H, Wang H, Shu Y, Li X. 2018. miR-103 Promotes Neurite Outgrowth and Suppresses Cells Apoptosis by Targeting Prostaglandin-Endoperoxide Synthase 2 in Cellular Models of Alzheimer’s Disease. Front Cell Neurosci 12:91.

65. Boscher E, Goupil C, Petry S, Keraudren R, Loiselle A, Planel E, Hébert SS. 2021. MicroRNA-138 Overexpression Alters Aβ42 Levels and Behavior in Wildtype Mice. Front Neurosci 14:591138.

66. Estfanous S, Daily KP, Eltobgy M, Deems NP, Anne MNK, Krause K, Badr A, Hamilton K, Carafice C, Amer AO et al. 2021. Elevated Expression of MiR-17 in Microglia of Alzheimer’s Disease Patients Abrogates Autophagy-Mediated Amyloid-β Degradation. Front Immunol 12:705581.

67. Liang C, Mu Y, Tian H, Wang D, Zhang S, Wang H, Liu Y, Di C. 2021. MicroRNA-140 silencing represses the incidence of Alzheimer’s disease. Neurosci Lett 758:135674.

